# An expression-directed linear mixed model (edLMM) discovering low-effect genetic variants

**DOI:** 10.1101/2023.07.13.548939

**Authors:** Qing Li, Jiayi Bian, Yanzhao Qian, Pathum Kossinna, Paul MK Gordon, Xiang Zhou, Xingyi Guo, Jun Yan, Jingjing Wu, Quan Long

## Abstract

Detecting genetic variants with low effect sizes using a moderate sample size is difficult, hindering downstream efforts to learn pathology and estimating heritability. In this work, by utilizing informative weights learned from training genetically predicted gene expression models, we formed an alternative approach to estimate the polygenic term in a linear mixed model (LMM). Our LMM estimates the genetic background by incorporating their relevance to gene expression. Our protocol, expression-directed linear mixed model (edLMM), enables the discovery of subtle signals of low-effect variants using moderate sample size. By applying edLMM to cohorts of around 5,000 individuals with either binary (WTCCC) or quantitative (NFBC1966) traits, we demonstrated its power gain at the low-effect end of the genetic etiology spectrum. In aggregate, the additional low-effect variants detected by edLMM substantially improved estimation of missing heritability. edLMM moves precision medicine forward by accurately detecting the contribution of low-effect genetic variants to human diseases.

## INTRODUCTION

Discovering genetic variants associated with risk of complex traits is a long-standing theme in the field of genetics. Given complicated genetic background of participating individuals, it is generally difficult to identify common variants that have statistically small effect sizes (e.g., with *R*^*2*^ <1%) using a moderate sample size (e.g., at the level of thousands)(Zondervan and Cardon 2004). Although their independent contributions are statistically weak in association studies, they may play critical roles in the pathology of individual patients, despite of their seemingly low contribution in the whole population. For instance, the Price lab has articulated that pathologically important variants have to explain low phenotypic variance due to the effect of negative selections(Gazal et al. 2018; O’Connor et al. 2019; Peyrot and Price 2021). Additionally, these low-effect variants might aggregately account for a substantial proportion of the heritability. If the above assumptions are indeed valid, methods discovering low-effect variants using a moderate sample size may uncover risk genetic variants in many applications, including the experimental investigation of pathology, formation of a polygenic risk score (PRS)(Choi et al. 2020), and characterization of the susceptibility genetic architecture of a complex trait.

Linear mixed models (LMMs) serve as primary approaches to discovering genetic variants associated with disease phenotype in association studies(Kang et al. 2008a; Listgarten et al. 2010; Price et al. 2010; Yu et al. 2006; Zhang et al. 2010). This is largely due to the advantage of using a random term modeled by the genetic relationship matrix (GRM), to correct population structure and other confounding effects. In this work, we determined that by accurately modeling this random term in an LMM with the informative weights learned from omics data, (e.g., transcriptome), one can discover risk genetic variants with lower effect size on diseases/traits, leading to more proportion of explainable heritability. From a technical standpoint, we extended an idea from transcriptome-wide associate studies (TWAS)(Gamazon et al. 2015; Gusev et al. 2016) using our own unique angle(Cao et al. 2022; Cao et al. 2021b). Here we provide an overview of our methodology, to understand what exactly LMM and TWAS are.

First, in its simplest form, the LMM can be generally expressed as: *y* ∼ *x* + *u* + *ε*, where *y* is the phenotype, *x* is the focal genetic variant under consideration, and *u*, the key component in LMM is the random term, which distinguishes LMM from a simple linear regression. Here *u* follows a multivariate normal distribution *MVN*(**0, *K***), where ***K***, the variance-covariance matrix of the MVN is estimated by the genome-wide variants(Powell et al. 2010; Rousset 2002; Speed and Balding 2015). ***K*** was initially called “kinship matrix” when pedigree data were used, and then called GRM when using seemingly unrelated individuals in a typical association study. Both names emphasize the accounting for “relatedness” among samples, which is the source of population stratification(Freedman et al. 2004; Kang et al. 2010b). Another interpretation of *u*, which is also popular in the field of genomic selection(Crossa et al. 2017; Xu 2013; Xu et al. 2021), is to call it a “polygenic” term(Goddard and Hayes 2007; Xie et al. 2021; Xu 2003; 2017), which captures the variance component explained by contributing variants (other than the focal variant *x*) in the genome. In this sense, the interpretation of “relatedness”, or “population structure” is a rough approximation of the “polygenic model” as the chance of two individuals carrying the same (combinations of) functional alleles may be proportional to their relatedness. Although the number of potential causal variants may be large, LMMs enjoy the advantage of not running into overfitting because the GRM is calculated in advance so that genetic variants with small contributions will not be modeled individually. However, LMMs may run into the risk of underfitting as the calculation of GRM evenly incorporates the genetic variants without explicitly considering their functional consequence. In this sense, modeling the GRM more accurately will increase the overall fit, leading to higher power in the association mapping(Fisher 1919; Ober et al. 2001; Visscher and Goddard 2019). Indeed, treating different variants selectively in GRM has been shown to improve performance in various applications, e.g., the LDAK model(Speed et al. 2017; Speed et al. 2012). However, incorporating informative weights in an LMM to better model its null distribution has not been explored yet. Specifically, although it is generally believed that the use of a polygenic term modeled by GRM could be equivalent to a form of Bayesian feature selection(Sorensen et al. 2002; Zhou et al. 2013) (a mathematical elaboration is enclosed in **Supplementary Notes I**), there is no such efforts to improve the null distribution modeling in LMM using functional *a priori*.

Second, we articulate our alternative understanding of what is TWAS (recently published in(Cao et al. 2021a; Cao et al. 2022; Cao et al. 2021b)). The mainstream format of TWAS is a two-step protocol: first, one forms an expression prediction model using (usually cis) genotype: ***e*** ∼ Σβ_i_***x***_***i***(*REF*)_ + ***ε*** in a reference dataset (e.g., GTEx(Carithers and Moore 2015; Lonsdale et al. 2013)), where *e* is the expression vector of the focal gene (subscript specific for the gene omitted for simplicity), ***x***_***i***(*REF*)_ are genotypes in the reference dataset, and β_i_ are the weights to be trained. The predicted expression is called Genetically Regulated eXpression, or GReX. Then, in the second step, in the main dataset for association mapping (that doesn’t contain expression values), one can predict expressions using the genotype: ê = Σ*β*_*i*_***x***_*i*(*GWAS*)_, where ê is the predicted expression β_i_ are the previously trained weights, and ***x***_*i*(*GWAS*)_ are the corresponding genotype in the GWAS dataset. Then using the predicted expression, one will conduct association mapping ***y*** ∼ê. Our recent works have shown that this standard interpretation of “predicting expressions” may need to be revisited. First, theoretical power analysis showed that TWAS could be more powerful than the hypothetical scenario in which expression data is available in the main GWAS dataset(Cao et al. 2021a); additionally, TWAS could be underpowered compared with GWAS when the expression heritability is low(Cao et al. 2021a). Both results question the interpretation of the prediction of expression in TWAS. As such, we proposed to interpret the “prediction” step as a selection of genetic variants directed by expression(Cao et al. 2022). From the perspective of Machine Learning, the first step in TWAS is exactly feature selection and the second being feature aggregation. With this interpretation in mind, one may cancel GReX and instead conduct feature selection and aggregation independently. Indeed, novel methods splitting these two steps developed by us(Cao et al. 2022; Cao et al. 2021b) and others(Tang et al. 2021) showed higher power than standard TWAS.

Based on our understanding of LMM and alternative interpretation of TWAS, we developed expression-directed linear mixed model, or edLMM, a synergy between LMM and TWAS, to model the polygenic term (based on a weighted GRM) using expression-based feature selection. In both real data and simulations, we showed that edLMM is more powerful than EMMAX(Kang et al. 2010b), a flagship LMM used by many researchers, and it is particularly sensible when the effect size is low. Additionally, it is demonstrated that the explainable proportion of heritability is substantially improved when the low-effect variants discovered by edLMM are incorporated.

## MATERIALS AND METHODS

### Overview of the edLMM model

edLMM is structured into two steps: first, like a typical TWAS protocol, BSLMM(Zhou et al. 2013) is employed to train a *cis* genotype-based expression prediction model using a reference panel such as GTEx(Carithers and Moore 2015; Lonsdale et al. 2013) (**Figure 1A**). Based on our alternative interpretation of TWAS, we do not have to use the predicted expression (GReX) in the next step; instead, the first step is used only for feature selection. We then pull together all the selected genetic variants (from all individual gene prediction models) and their BSLMM-coefficients to form the GRM. Instead of the traditional GRM formation using the product of a centralized genotype matrix and its transpose, we incorporate BSLMM-trained coefficients as functional weights by inserting a diagonal matrix to rescale the product (**Figure 1B**). This GRM weighted by expression-relevance will be then used in a standard LMM (**Figure 1C**). Note that, the use of weighted GRM is equivalent to adding functional *a priori* in a Bayesian setting (**Supplementary Notes I**). The edLMM protocol enjoys higher power for discovering low-effect variants due to its more accurate estimation under the null model: when the effect size is small, the subtle distance between the actual and the modeled null distributions may be larger than the effect size, masking the genuine signal to be captured (**Figure 1D**). In contrast, by better modeling the null using the weighted GRM, edLMM effectively amplified the statistical signal to alleviate this difficulty (**Figure 1E**). As such, the same effect size in the fixed term (i.e., the regression coefficient) will be assessed to a more significant p-value in edLMM, increasing statistical power.

**Figure 1.**
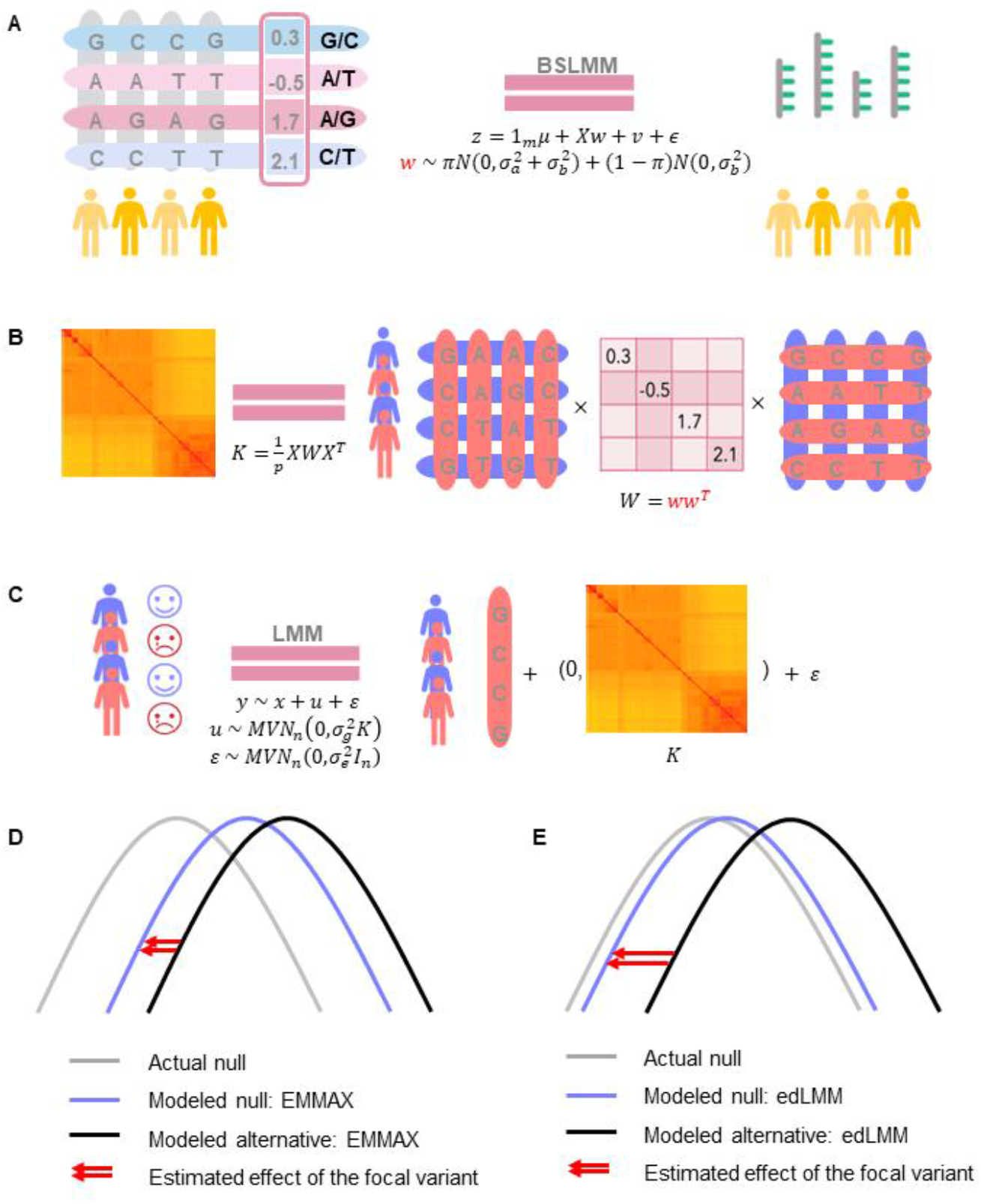
The edLMM protocol. **A)** The BSLMM tool (Bayesian sparse linear mixed model) is used to train the weights of each genetic variant, using genotype-expression data from a reference panel such as GTEx. **B)** The calculated weights are used to compute the weighted GRM in a GWAS dataset. **C)** The weighted GRM is used as the variance-covariance matrix of the multivariate normal distribution in an LMM for the GWAS. **D)** and **E)** show the null and alternative distributions of genetic variants under EMMAX (the standard LMM) and edLMM protocol. The newly formed null distribution in (e) is more accurate, signifying the statistical effect of the focal variant (red arrows).

### Mathematical formulation of the standard LMM and edLMM

The standard linear mixed model in genotype-phenotype association mapping is defined as:

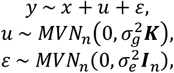

where *n* is the number of individuals, *y* is an *n* × 1 vector of quantitative or qualitative phenotypes, *x* is an *n* × 1 vector of genotype, *u* is the random effect term with 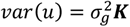 where ***K*** is the *n* × *n* genetic relationship matrix (GRM), *ε* is an *n* × 1 vector of residual effects such that 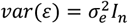 where *I*_*n*_ is an *n* × *n* identity matrix, *MVN*_*n*_ denotes multivariate normal distribution with *n* dimension. The overall phenotypic variance-covariance matrix can be expressed as 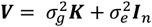. (**Figure 1C**)

In standard LMM(Kang et al. 2010a; Kang et al. 2008b), the GRM ***K*** was calculated without assuming any functional weights, e.g., using the realized relationship matrix (RRM):

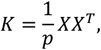

where *X* is the *n* × *p* standardized genotype matrix, and *p* is the number of genetic variants.

In edLMM, we calculated ***K*** by weighting the genetic variants according to their contributions to gene expressions. Specifically, the weighted realized relationship matrix (w-RRM) is calculated as (**Figure 1B**):

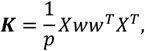

where *w* is a *p* × 1 vector of weights estimated from BSLMM, a Bayesian sparse linear mixed model that predicts expressions based on genotype(Zhou et al. 2013): (**Figure 1A**):

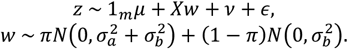

Here *z* is a vector of gene expressions measured on *m* individuals, *m* is the number of individuals in genotype-expression data set, 1_*m*_ is an *m* × 1 vector of 1*s, μ* is a scalar representing the phenotype mean, *X* is an *m* × *p* matrix of genotypes measured on the same individuals at *p* genetic variants, *w* is the corresponding effects of variants which come from a mixture of two normal distributions, *v* is the random effect term, and *ϵ* is the error term. In the mixture normal distribution, *π* controls the proportion of *w* values that are non-zero. 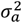 controls the expected magnitude of the non-zero *w*. 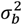 controls the expected magnitude of the random effects *v*. The BSLMM model involves *μ, π*, 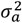 and 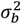 as hyperparameters and specifies their prior distributions. *w* can be estimated using Markov chain Monte Carlo (MCMC) given the observed data.

### GWAS and gene expression datasets, genotype imputation and quality controls

To test the performance of edLMM in real data, two representative GWAS datasets with around 5,000 samples for binary and quantitative traits, namely WTCCC (The Wellcome Trust Case Control Consortium)(Wellcome Trust Case Control 2007) and NFBC1966 (Northern Finland Birth Cohort 1966)(Sabatti et al. 2009) were used. WTCCC contains ∼2,000 individuals for each of the seven diseases and a shared set of ∼3,000 controls: type 2 diabetes (T2D), type 1 diabetes (T1D), hypertension (HT), rheumatoid arthritis (RA), coronary artery disease (CAD), Crohn’s disease (CD) and bipolar disorder (BD) (**Supplementary Table S1**). We removed genetic variants with a minor allele frequency (MAF) <= 1%, resulting in a pruned set of 392,937 variants carried forward in our analysis. NFBC1966 contains quantitative metabolic traits: triglycerides, high-density lipoprotein (HDL), low-density lipoprotein (LDL), glucose, insulin, C-reactive protein, body mass index, and systolic and diastolic blood pressure (**Supplementary Table S2**). A full set of 299,681 variants remained after removing variants with MAF <= 1%, along with 5,244 individuals. A variant will be reported significant if it reaches the Bonferroni corrected significance, i.e., 0.05 divided by number of genome-wide genetic variants in the dataset.

Replication studies for significant variants were conducted using two independent dbGaP datasets for T1D(Pezzolesi et al. 2009) (dbGaP: phs000018.v2.p1; N =1,792 #SNP =364,292) and T2D(Wolf et al. 2003) (phs000237.v1.p1, N =1,384, #SNP =1,199,187) respectively. The genomic build for these two datasets was hg18 causing few overlapped SNPs with WTCCC after they were lifted over to hg38. Therefore, we conducted imputation to increase SNPs number in a secure manner. Specifically, genotype imputation was conducted using a pipeline involving SHAPEIT4(Delaneau et al. 2019) for pre-phasing and IMPUTE5(Rubinacci et al. 2020). Datasets aligned to previous versions of the human genome were lifted over to hg38 using CrossMap(Zhao et al. 2014) and SHAPEIT4 was used with the included maps for hg38 to produce pre-phased VCFs of each chromosome with no missing genotypes. After that, IMPUTE5 was used to output chunked VCFs each containing imputed genotypes using data from the 1000 Genome Project resulting in over 73 million SNPs for both final datasets. Finally, imputed SNPs with the same genomic coordinates as WTCCC (392,937) SNPs were selected for downstream EMMAX and edLMM analyses (390,838 for T1D and 389,577 for T2D). A genetic variant is reported as a being successfully replicated if it meets two criteria: 1) its p-value is lower than 0.05 divided by number of significant SNPs in corresponding WTCCC disease dataset; and 2) it locates within 500K base-pair distance to the WTCCC significant variant to be replicated.

Gene expression data generated by RNA-seq were obtained from the Genotype-Tissue Expression Project version 8 (GTEx v8)(Consortium 2015). As the sample size and data quality vary among different GTEx tissues, the number of genes for each tissue also varies. Here we used Whole-blood sample (N = 670) as it is the most representative general dataset to use when the disease-causing tissue is unclear. The target tissue has a total of 20,315 genes. As the focus of edLMM is to collect a large number of genetic variants to form the GRM, in contrast to make an accurate gene prediction model that conventional TWAS do, we did not conduct filtering of genes and used all 20,315 genes. The covariates, including genotyping principal components (PCs), were obtained from GTEx_Analysis_v8_eQTL_covariates.tar.gz, which was downloaded from the GTEx portal. For each gene, the gene expression was adjusted for top five genotyping PCs, age, sex, sequencing platform, PCR protocol, and 60 peer confounding factors using PEER, following the GTEx best practices(Stegle et al. 2012).The BSLMM training used GTEx genotype within 1Mb flanking region of the focal genes.

### Simulation study

We conducted simulations to estimate power and Type-I Error of edLMM, in contrast to EMMAX. The genotype data of NFBC (N = 5,244) and whole-blood gene expression data of GTEx (v8, N = 670) were used to simulate phenotypic values. Quantitative traits and binary traits were simulated separately, which are described below.

Quantitative phenotypes were simulated based on an “additive” architecture, combing the contributions from a focal variant, polygenic background, and a noise. During each round of the simulation, we first simulated one variant as the target to be tested by the models (i.e., EMMAX and edLMM). Then we randomly sampled 500 genes from a total of 20,315 in GTEx v8. For each gene, two-cis genetic variants were randomly selected. These 500 × 2 = 1,000 variants will form the contribution of polygenic term. The noise was simulated by rescaling the phenotypic variance components to ensure that the phenotypes have different pre-specified heritability. Finally, phenotypes were calculated by adding the contribution from the focal variant, weighted sum of 1,000 variants from GTEx v8 where weights were from the pre-trained BSLMM model (polygenic background), and the noise. We repeated simulations 1,000 times for both quantitative and qualitative phenotypes.

Mathematically, the model is given by:

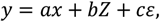

where *x* represents the focal variant, 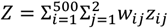, is the weighted sum of 1,000 variants with *w*_*ij*_′*s* pre-specified by the BSLMM model, *ε* is the noise sampling from a standard normal distribution (*ε* ∼ *N*(0,1)). The coefficients *a, b, c* are calculated as follows.

The overall heritability (i.e., variance components contributed by all variants, *x* and *Z*) is fixed to 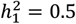. The variance component of the focal variant, 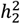, is set to be 9 different quantities: 0.001, 0.002, 0.003, 0.004, 0.005, 0.006, 0.007, 0.01 and 0.015. The variance component of noise term is 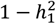. To ensure 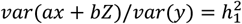, we calculate *a, b, c* as follows. We first set *a* = 1. Given *var*(*ε*) = 1 *b, c* can be solved using the equations below:

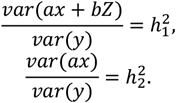

Solving the above equations yields:

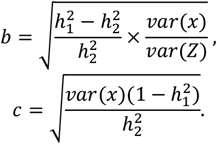

The model for simulating binary traits largely reused the above model for quantitative traits. We first simulated a quantitative trait, and then used a liability cut-off to divide data into cases and controls, keeping the ratio between cases and controls roughly 2:3, mirroring the WTCCC dataset for which we have conducted the real data analysis.

### Type-I error estimation and power calculations

The Type-I error of each protocol was estimated around *α* = 0.05. To achieve this, for the binary case, we randomly assigned phenotypes (1 for a case and 2 for a control) to 5,244 subjects to empirically determine the null distribution of each protocol. In the quantitative case, we randomly assigned phenotypes by drawing samples from a standard normal distribution. Then we ranked p-values calculated for each variant and took the top 5% as the cut-off. If this cut-off is around 0.05, then it indicates the Type-I error is under control.

For the above genetic architecture and its associated parameters, 1,000 datasets containing simulated phenotypes were created. In each simulated data set, we tested the two protocols’ (EMMAX and edLMM) ability to successfully identify the causal variant simulated from GWAS data set. Success is defined as the Bonferroni-corrected p-value of this variant that is lower than a predetermined critical value (0.05). We then calculated the power of each protocol as the number of successes divided by the number of rounds of simulation (1,000).

### Protocols for association mapping using edLMM and EMMAX

First, BSLMM(Zhou et al. 2013), a function provided by the GEMMA toolkit (Zhou and Stephens 2012), was applied to the GTEx (v8) dataset to train the weights of each *cis* genetic variant (in the surrounding 1Mb flanking region). Using the default setting of GEMMA, the variant effect *w* can be decomposed into two parts: *α* that captures the minor effects that all variants have, and *β* that captures the additional effects of some large effect variants. The total effect size for a given genetic variant *i* is then *w*_*i*_ = *α*_*i*_ + *β*_*i*_. Based on the formulas described in “*Mathematical formulation of the standard LMM and edLMM*” above, we implemented a new function, ‘kinshipwe’, in our tool Jawamix5(Xiong et al. 2019) (Long et al. 2013) to calculate the weighted GRM. An existing function, ‘kinship’, is used to calculate the standard (unweighted) GRM. Then, taking the above two different versions of GRM as alternative input (representing edLMM and EMMAX, respectively), the ‘emmax’ function of Jawamix5 was used to conduct the GWAS for both WTCCC and NFBC.

### SNPs annotated to genes

Significant SNPs identified by edLMM and EMMAX were annotated to genes using cS2G (Combining SNP-to-gene) (Gazal et al. 2022), which utilized multiple combined SNP-to-gene strategies and trained on various diseases.

### Genes’ functional validations

To verify functional relevance of statistical results out of the association mapping, we performed gene-disease association analysis using the DisGeNET repository (v7.0)(Pinero et al. 2017; Pinero et al. 2015), a platform containing 1,134,942 gene-disease associations (GDAs), between 21,671 genes and 30,170 traits. For each disease, DisGeNET has its specific summary of gene-disease associations containing information of identified significant genes and their gene-disease associations scores (GDAs). This summary of associations can be searched using the CUI number of each disease. The CUI numbers of seven diseases in the WTCCC are listed **Supplementary Table S3**. The gene model file used in this study was downloaded from the GTEx website (https://www.gtexportal.org/home/datasets/gencode.v26.GRCh38.genes.gtf).

### Heritability estimation

We aim to quantify the degree to which the low-effect variants bring incremental heritability. We estimated SNP-heritability identified by both protocols in the WTCCC and NFBC datasets using GCTA(Yang et al. 2011) (Genome-wide Complex Trait Analysis), a software package developed to estimate the proportion of phenotypic variance explained by all genome-wide variants. In particular, **GCTA-GRM** was used to calculate the genetic relationship matrix (GRM) from the associated variants and **GCTA-GREML** was used to estimate the variance explained by them.

## RESULTS

### Simulations confirm the higher power of edLMM at the low-effect size end of the genetic effects spectrum

We conducted simulations to test the statistical power of edLMM against EMMAX(Kang et al. 2010b), based on both quantitative and binary traits (**Materials and Methods**). First, the type-I errors in both binary and quantitative datasets were around 0.05, which are well controlled (**Supplementary Table S3**). The power comparisons showed that, for both quantitative traits and binary traits (=case/control), edLMM outperforms EMMAX by a substantial margin when the variance component explained by the focal genetic variant is low, i.e., less than 0.01 **(Figure 2A and 2B; Supplementary Table S4)**. In contrast, when the variance component is high, e.g., above 0.01, there is no clear distinction between the two protocols. This supports our assertion that by fitting random terms using weighted GRM in LMMs, the statistical power is increased for variants with low effect sizes.

**Figure 2:**
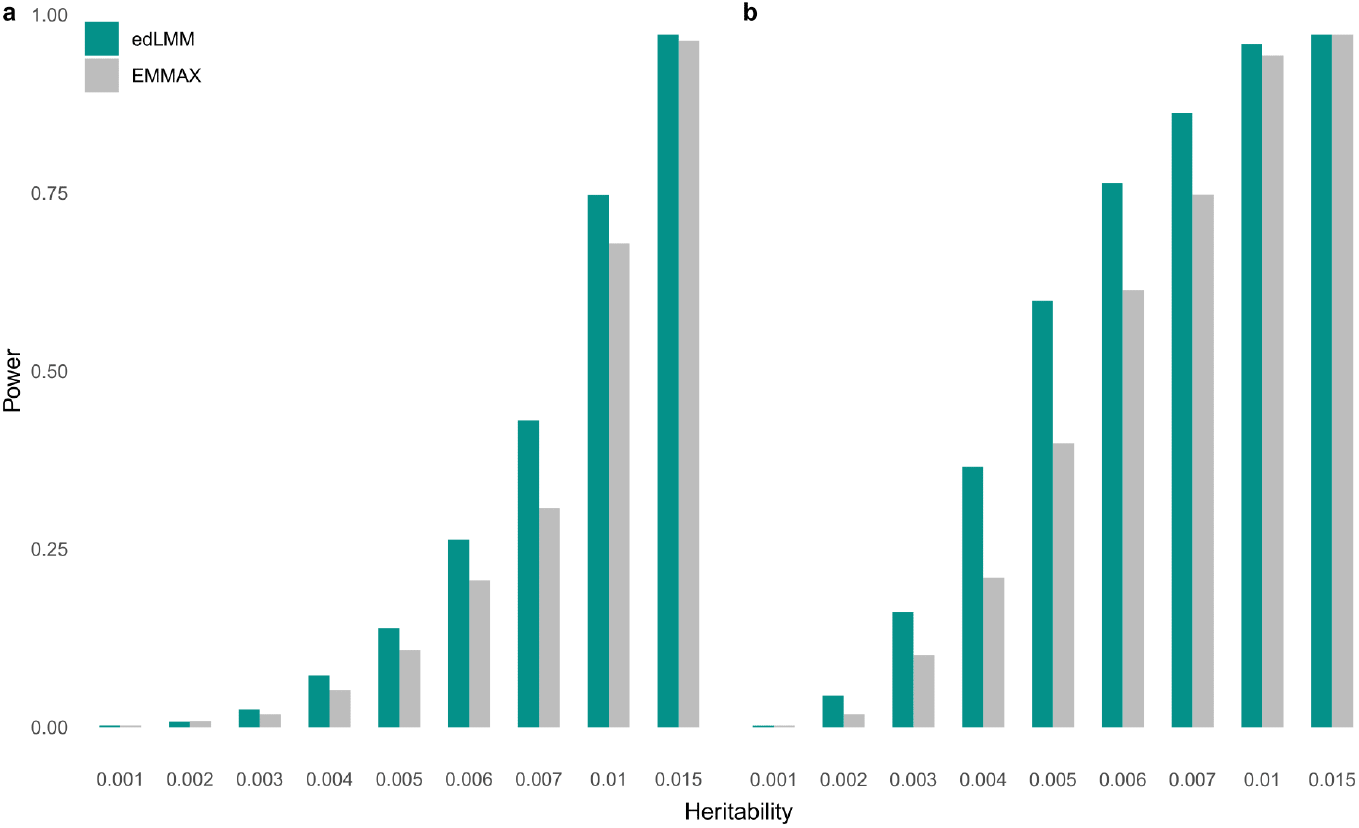
Power comparison between edLMM and EMMAX based on simulations. **A)** binary and **B)** quantitative traits. The variance component explained by the focal variant, 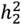, which is depicted in the x-axis, is set to 9 different quantities: 0.001, 0.002, 0.003, 0.004, 0.005, 0.006, 0.007, 0.01 and 0.015.

### edLMM improves identification of genetic variants in real data

We applied edLMM and EMMAX to two frequently used benchmark datasets that have a moderate sample size of around 5,000 (**Materials and Methods**). Excluding the traits for which none of the protocols called any signal (after Bonferroni correction), edLMM identified more significant variants than EMMAX in both WTCCC (**Figure 3; Supplementary Table S5)** and NFBC (**Supplementary Table S6**). In addition, we replicated EMMAX and edLMM significant discoveries in two independent datasets for T2D and T1D using other independent GWAS datasets (**Materials and Methods**). Our results showed that edLMM is more robust than EMMAX in both diseases, as edLMM replication rates are 11.97% (14/117) and 10.16% (31/305) in T2D and T1D, respectively, in contrast to EMMAX’s replication rates of 10.53% (6/57) and 4.97% (9/181).

**Figure 3:**
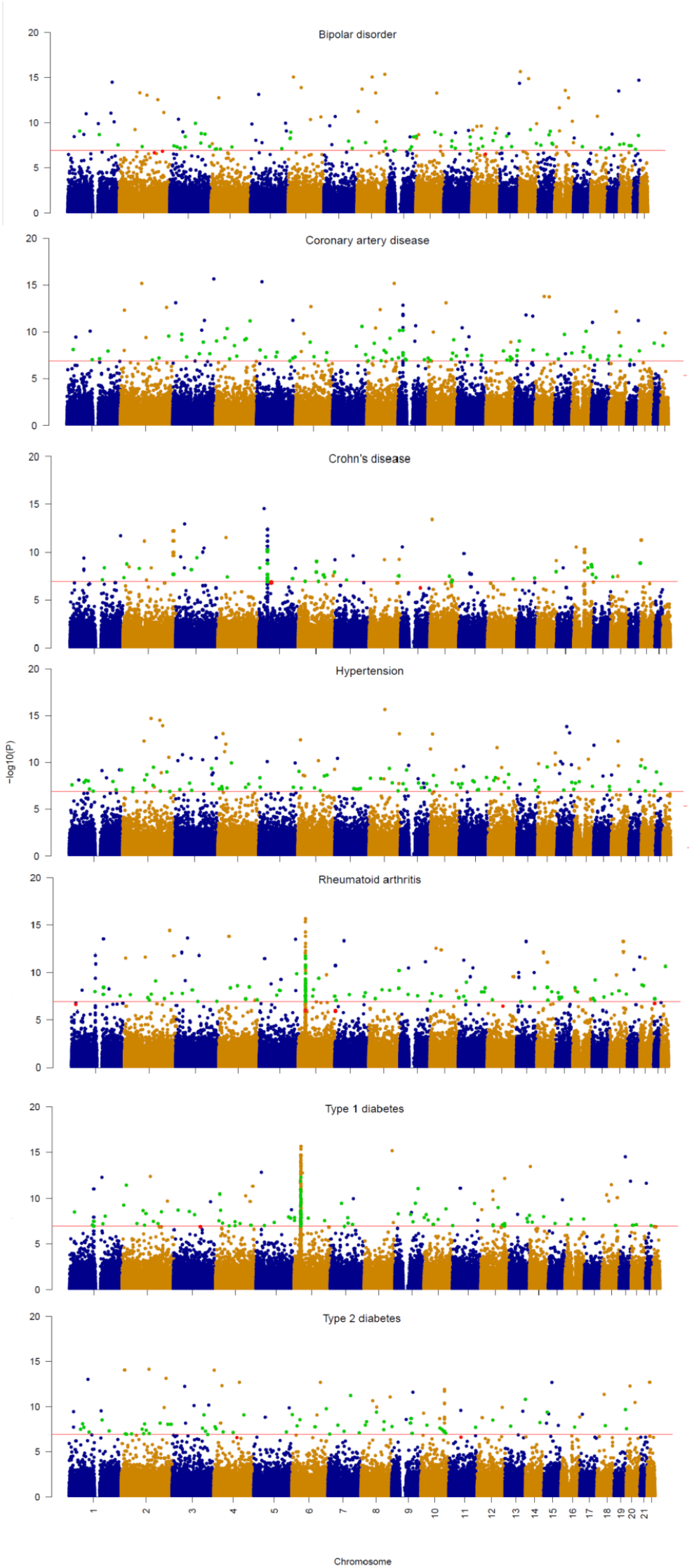
Manhattan plots of significant genetic variants identified by edLMM of the seven WTCCC diseases. Unique significant SNPs identified by edLMM are colored in Green and the unique ones for EMMAX are colored in Red. Red vertical lines stand for the cut-off of Bonferroni correction.

### edLMM discovered additional genetic variants with low effect sizes

We further analyzed whether edLMM’s superior performance over EMMAX is indeed due to the advantage of detecting low-effect variants (that is indicated by our design intuition and simulations). To this end, the distribution of effect sizes (= absolute values of regression coefficient of the genetic variant) of significant signals identified by EMMAX and edLMM were visualized for WTCCC (**Figure 4**). It is observed that edLMM outperforms EMMAX across the spectrum, and the advantage for low-effect variants (<=0.05) are particularly pronounced.

**Figure 4:**
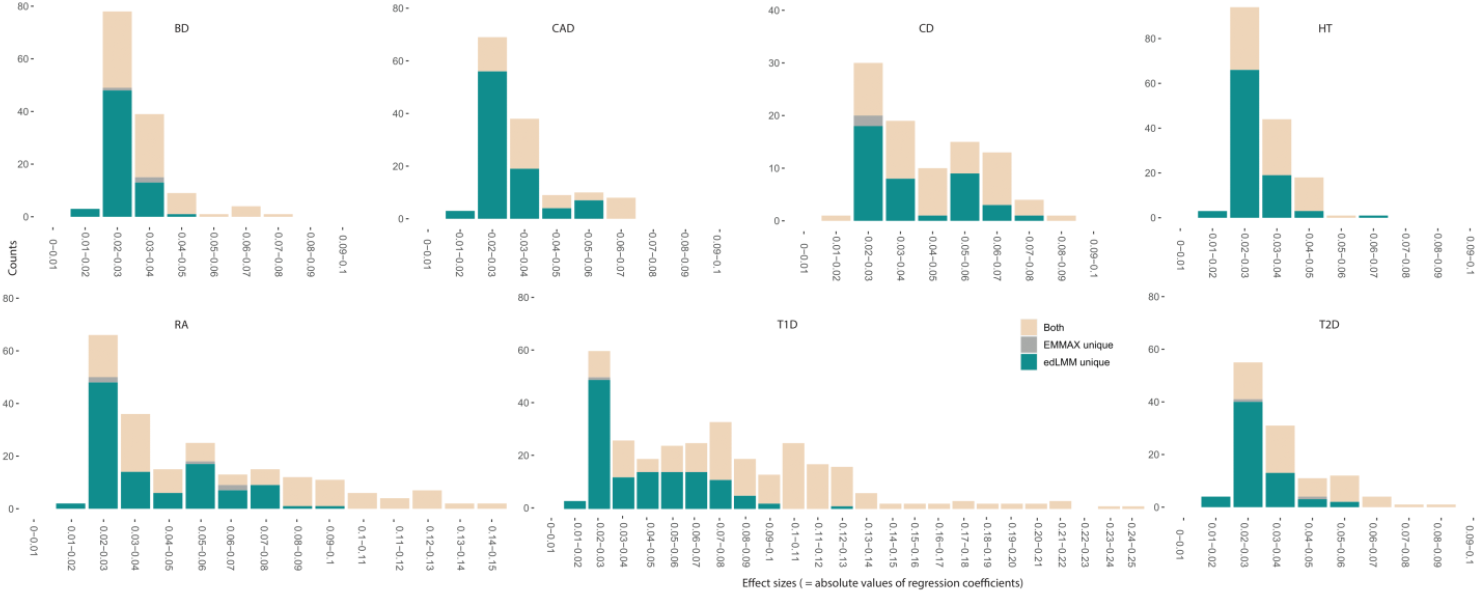
Distribution of effect sizes of significant genetic variants identified by edLMM and EMMAX of the seven WTCCC diseases. Variants identified by both EMMAX and edLMM (Yellow), uniquely identified by edLMM (Green) and uniquely identified by EMMAX (Grey) are depicted.

### edLMM-identified variants are functionally relevant

To check the functional relevance of the statistical results, we first mapped each significant variant to genes using cS2G, a state-of-the-art tool to map genetic variants to genes based on various factors(Gazal et al. 2022). Then we queried the role of genes in the related disease in DisGeNET, an established repository of validated genes associated with diseases(Pinero et al. 2017) (**Materials and Methods; Supplementary Table S7**). We compared edLMM and EMMAX for each disease on their numbers of validated and the rate of validated (number of validated divided by the total number of discovered). Additionally, by removing the variants that were discovered by both methods, we also calculated the number of unique validated as well as unique validated rate. We find that edLMM outperformed EMMAX in all the seven WTCCC diseases in terms of numbers of validated, although validated rates are slightly lower (**Figure 5; Supplementary Table S7**).

**Figure 5:**
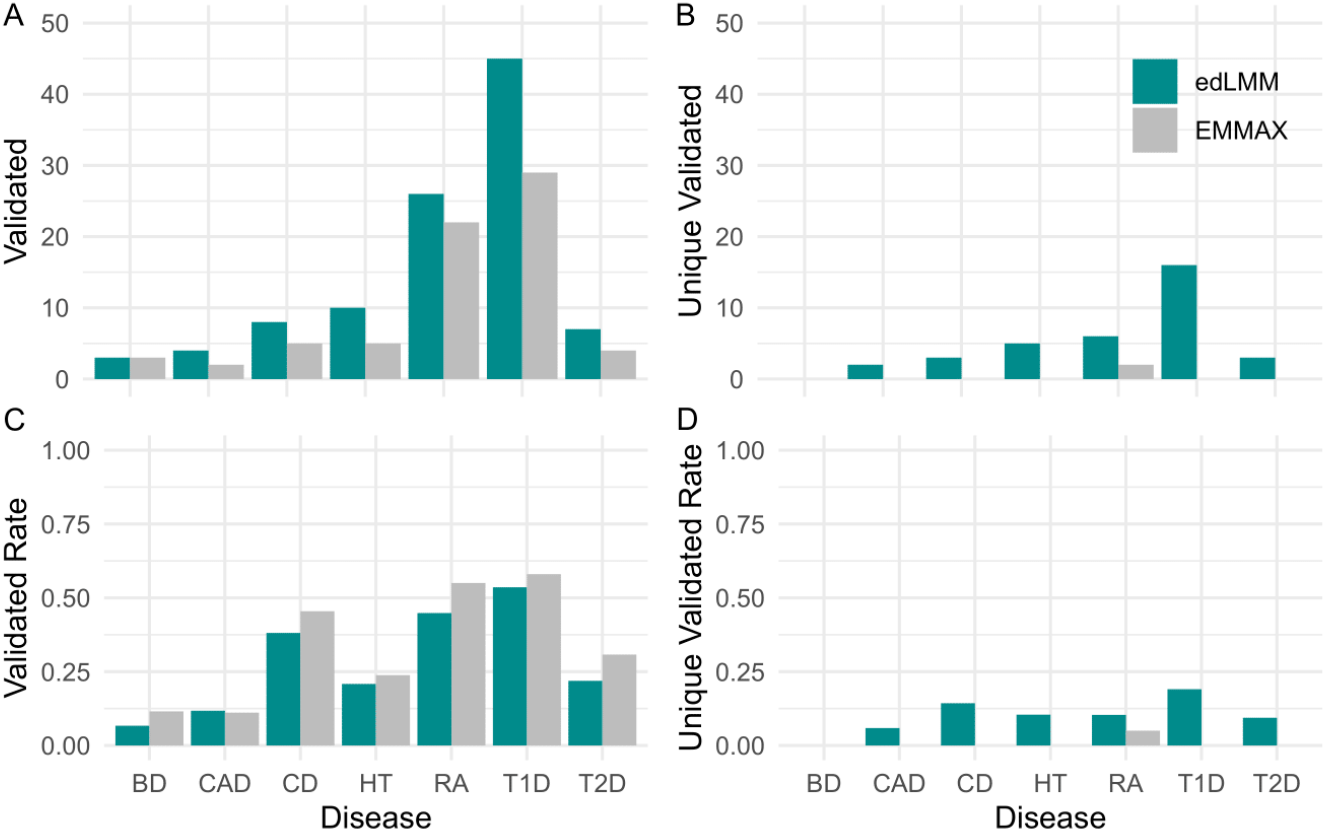
The DisGeNET functional annotations of signals identified by edLMM and EMMAX in the seven WTCCC diseases. **A**) Number of validated genes (validated) reported in the DisGeNET database. **B**) Number of validated genes (unique validated) reported exclusively by one of the two protocols in the DisGeNET database. **C**) Proportion (validated rate) of all reported genes which are validated in the DisGeNET database. **D**) Proportion (unique validated rate) of genes reported exclusively by one of the two protocols in the DisGeNET database.

However, the edLMM validated rates are still good considering the substantially larger number of candidates and their low effect sizes, as well as the fact that low-effect variants may be less studied and deposited in the databases. All significant genes identified by any of the two protocols for the seven diseases of WTCCC, along with their p-values, effect size and MAF, are in **Supplementary Tables S8-14**. For NFBC data set, we focused on two traits: high-density lipoprotein (HDL) and low-density lipoprotein (LDL), as other traits the number of significant variants identified by EMMAX and edLMM is relatively small and not much difference to analyze. Following the same protocol, we mapped significant variants identified by both EMMAX and edLMM to genes and reported these genes’ DisGeNET validations as well (**Supplementary Tables S15, S16**).

### Literature support of genes uniquely identified by edLMM

We conducted a literature search on the genes that were discovered by edLMM but not EMMAX. For WTCCC, we focused on T1D and RA, which contain most significant genes among seven diseases. For each disease, we selected two edLMM uniquely discovered genes based on their DisGeNET scores, as well as two genes that are not reported by DisGeNET but with large cS2G scores. For NFBC, only one gene from HDL was annotated by cS2G but has not been reported by DisGeNET. The gene names and associated p-values and scores are presented in (**Supplementary Tables S17**). We were able to identify multiple lines of evidence in literature for these genes (**Supplementary Notes II**).

### Inclusion of edLMM-identified genetic variants leads to higher explainable SNP-heritability

To quantify the incremental benefits brought by low-effect variants, we collected variants identified by edLMM and EMMAX to estimate SNP heritability using GCTA, a standard tool for heritability estimation(Yang et al. 2011) (**Materials and Methods**). In the WTCCC dataset, the SNP heritability explained by associated variants increased substantially, ranging from 6.76% (T1D) to 147.61% (HT), with the median being 73.29% (T2D) (**Figure 6; Supplementary Table S18**). In the NFBC dataset, SNPs’ heritability increases of 24.44% for HDL and 1.35% for LDL were observed (**Supplementary Table S19)**.

**Figure 6:**
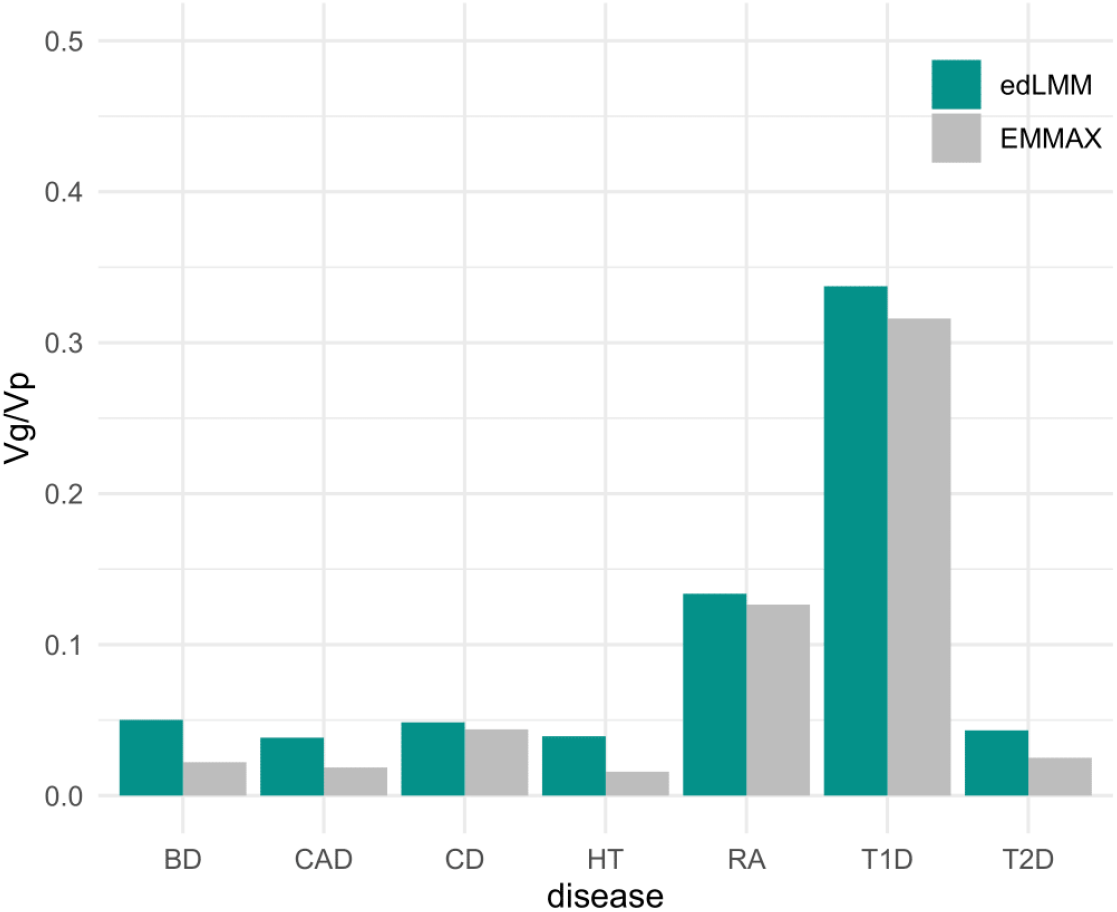
The proportion of phenotypic variance explained by significant genetic variants (Vg/Vp) identified by edLMM and EMMAX in the WTCCC dataset.

## DISCUSSION

Based on our interpretation of LMM and unique understanding of TWAS(Cao et al. 2022; Cao et al. 2021b), we proposed edLMM, a new model that borrows a variant-weighting protocol to refine the modeling of the polygenic term (represented by the weighted GRM) in an LMM model. This design has been shown to bring a substantial power gain when detecting low-effect genetic variants in association studies with moderate samples (i.e., WTCCC and NFBC that have around 5,000 individuals), leading to improved SNP-heritability estimations. This work further suggests that, by skipping the use of GReX, our disentangling of genetic feature selection and aggregation in transcriptome-wide association studies(Cao et al. 2022) may open a door for novel models integrating multi-scale -omics to be developed.

This work has been only focusing on expression data within a single tissue. Natural extensions include developing refined protocols using multi-tissue-based variants selection(Hu et al. 2019) and the inclusion of other functional -omics data such as DNA methylation(Wu et al. 2020), CHIP-Seq(He et al. 2022), etc. The flexibility of the edLMM framework enables straightforward incorporation of extra -omics data and multiple tissue-based feature selections, which are our future work. Another extension is to use GLMM to handle binary traits, although it has been pointed out that ordinary LMM is also generally appropriate(Kang et al. 2010b).

Although there are many interpretations for “missing heritability” including rare variants(Zuk et al. 2012) and nonlinear interactions(Zuk et al. 2014), here we chose to focus on genetic variants with a low effect size. These alternative interpretations do not have to be mutually exclusive. Indeed, although our additional low-effect variants can explain substantial missing heritability, still most of the heritability is left unexplained. However, our contribution from the perspective of the linear contribution of low-effect variants may be the most straightforward to implement in practical genetic tests using PRS, as it incorporates common variants in a simple way.

## Funding

New Frontiers in Research Fund and an HBI pilot grant NFRFE-2018-00748 (Q.L.)

Alberta Innovates LevMax-Health Program Bridge Funds 222300769 (Q.L.)

Canada Foundation for Innovation 36605 (Q.L.)

NSERC Discovery Grant (RGPIN-2018-04328 J.W.; RGPIN-2017-04860: Q.L.)

Campbell McLaurin Chair for Hearing Deficiencies (J.Y.)

Alberta Innovates Graduate Student Scholarships (J.B.)

## Author contributions

(Q.L.1= Qing Li and Q.L.2 = Quan Long) Conceived the project: J.W. and Q.L.2; Supervised the study: J.W. and Q.L.2; Designed statistical model and derived mathematical formulations: J.B. & Q.L.2; Implemented computational tool: Q.L.1, J.B. and Y.Q.; Conducted analyses: Q.L.1, J.B., P.K.; Provided advice and tools: P.G., X.Z., X.G., and J.Y.; Provided funding support: J.Y., J.W., Q.L.2; Wrote the manuscript: Q.L.1, J.B., and Q.L.2, with contributions from all co-authors.

## Data and materials availability

GTEx gene expression: https://gtexportal.org/home/datasets

GTEx whole genome sequencing data: https://www.ncbi.nlm.nih.gov/projects/gap/cgi-bin/study.cgi?study_id=phs000424.v9.p2

WTCCC genotype: https://www.wtccc.org.uk/

NFBC genotype: https://www.ncbi.nlm.nih.gov/projects/gap/cgi-bin/study.cgi?study_id=phs000276.v2.p1

T1D and T2D dataset used in replication: https://www.ncbi.nlm.nih.gov/projects/gap/cgi-bin/study.cgi?study_id=phs000018.v2.p1 and https://www.ncbi.nlm.nih.gov/projects/gap/cgi-bin/study.cgi?study_id=phs000237.v1.p1

The gene model file: https://www.gtexportal.org/home/datasets/gencode.v26.GRCh38.genes.gtf

edLMM is a function in Jawamix5, which is publicly available at: https://github.com/theLongLab/Jawamix5

cS2G code is publicly available at: https://alkesgroup.broadinstitute.org/cS2G/code

## Supplementary Tables

**Supplementary Table S1**. Estimated empirical type-I errors of two protocols, EMMAX and edLMM, in two GWAS datasets.

**Supplementary Table S2**. Power comparison between EMMAX and edLMM based on simulation using binary traits and quantitative traits.

**Supplementary Table S3**. Description of the seven diseases in the WTCCC dataset.

**Supplementary Table S4**. Description of the nine phenotypes in the NFBC dataset.

**Supplementary Table S5**. Number of significant genetic variants identified by the two protocols for the seven diseases in the WTCCC dataset.

**Supplementary Table S6**. Number of significant genetic variants identified by the two protocols for the nine traits in the NFBC dataset.

**Supplementary Table S7**. The DisGeNET functional annotations of the seven diseases in the WTCCC dataset.

**Supplementary Table S8**. All significant genes identified by cS2G in the WTCCC T2D.

**Supplementary Table S9**. All significant genes identified by cS2G in the WTCCC T1D.

**Supplementary Table S10**. All significant genes identified by cS2G in the WTCCC HT.

**Supplementary Table S11**. All significant genes identified by cS2G in the WTCCC RA.

**Supplementary Table S12**. All significant genes identified by cS2G in the WTCCC CAD.

**Supplementary Table S13**. All significant genes identified by cS2G in the WTCCC CD.

**Supplementary Table S14**. All significant genes identified by cS2G in the WTCCC BD.

**Supplementary Table S15**. All significant genes identified by cS2G in the NFBC HDL.

**Supplementary Table S16**. All significant genes identified by cS2G in the NFBC LDL.

**Supplementary Table S17**. Description of the genes for literature search in the WTCCC and NFBC datasets.

**Supplementary Table S18**. The proportion of phenotype variance explained by significant genetic variants (Vg/Vp) of the two protocols in the WTCCC dataset calculated by GCTA.

**Supplementary Table S19**. The proportion of phenotype variance explained by significant genetic variants (Vg/Vp) of the two protocols in the NFBC dataset calculated by GCTA.

