## Supplementary Files for "An expression-directed linear mixed model (edLMM) discovering low-effect genetic variants"

**Supplementary Notes I:**

**The equivalence between an LMM and a Bayesian feature selection (BSLMM), and the role of functional weights in edLMM**

A linear mixed model (LMM) is typically formulized as:

$$y \sim x + u + \varepsilon$$

where, $y$ is the vector of centralized phenotype (removing the intercept from the model), $x$ is the fixed effect representing the focal genetic variant, and $u$ is the random effect term with $var(u) = \sigma_g^2 \boldsymbol{K}$, $\varepsilon$ is the residual with $var(\varepsilon) = \sigma_e^2 \boldsymbol{I}$.

If we calculate $\boldsymbol{K}$ by standard GRM method, $\boldsymbol{K} = \boldsymbol{X}\boldsymbol{X}^T/p$, where $\boldsymbol{X}$ is the genome-wide genotype matrix and $p$ is the number of genome-wide variants, then we can prove the equivalence between the above LMM and Bayesian feature selection adapted by BSLMM as follows.

In LMM, under the null hypothesis when the effect of $x$ is zero, denoting $\Phi(\boldsymbol{\mu}, \boldsymbol{\Sigma})$ as the density of a muti-variate normal distribution with $\boldsymbol{\mu}$ as the mean vector and $\boldsymbol{\Sigma}$ as the variance-covariance matrix, the likelihood function is:

$$L\left(y \middle| \boldsymbol{X}, \sigma_g^2, \sigma_e^2\right) = \Phi\left(0, \sigma_g^2 \boldsymbol{K} + \sigma_e^2 \boldsymbol{I}\right)$$

Assuming only the genetic component is the focus, then we have $y_g$ the genetic contribution of $y$ can be expressed as:

$$L\left(y_g \middle| X, \sigma_g^2\right) = \Phi\left(0, \sigma_g^2 K\right) = \Phi\left(0, \sigma_g^2 X X^T/p\right)$$

In the Bayesian selection model, in particular BSLMM, we assume that:

$$y = 1_n \mu + \boldsymbol{X}\beta + \varepsilon$$

Where $\boldsymbol{X}$ is again the same genotype matrix mentioned above, and $\beta$ is the vector of effects corresponding to each genetic variant. Note that here the intercept $\mu = 0$ as the $y$ is centralized. Using the trivial priori in which all genetic variants are treated equally, we have that for all $i$, the $i$_th element of vector $\beta$, denoted as $\beta_i$, follows the same distribution:

$$\beta_i \sim N(0, \sigma_g^2/p)$$

Therefore, the variance-covariance matrix of the vector $y_g = X\beta$ can be derived by:

$$var(\boldsymbol{X}\beta) = \boldsymbol{X} var(\beta) \boldsymbol{X}^T = \boldsymbol{X}(\sigma_g^2/p)\boldsymbol{X}^T = \sigma_g^2 \boldsymbol{X}\boldsymbol{X}^T/p = \sigma_g^2 \boldsymbol{K}$$

In our new framework, edLMM, the improvement of adding functional weights may be explained in a Bayesian perspective as below:

If one aims to improve the priori distribution of $\beta$ for each SNP by adjusting the priori distributions of different genetic variants differently:

$$\beta_i \sim N(0, w_i^2 \sigma_g^2 / p)$$

Then the effect of above alternative functionally weighted priori is equivalent to an LMM by rescaling the GRM $\boldsymbol{K}$ by $\boldsymbol{w} = diag(|w_1|, |w_2|, \dots, |w_p|)$. To see why this is the case, denoting:

$$\hat{\beta} = \boldsymbol{w}\beta = diag(|w_1|, |w_2|, \dots, |w_p|)\beta$$

as the new vector of coefficients, we have

$$var(\boldsymbol{X}\hat{\beta}) = \boldsymbol{X}var(\hat{\beta})\boldsymbol{X}^T = \boldsymbol{X}\boldsymbol{w}(\sigma_g^2/p)\boldsymbol{w}^T\boldsymbol{X}^T = \sigma_g^2\boldsymbol{X}\boldsymbol{w}\boldsymbol{w}^T\boldsymbol{X}^T/p = \sigma_g^2\boldsymbol{K}_{\boldsymbol{w}}$$

As such, we conclude that the LMM is equivalent to a Bayesian feature selection model with trivial priori; and edLMM, which rescaled the GRM by functional weights, is equivalent to adding functional eights to the priori distributions.

**Supplementary Notes II:**

### Literature support of discoveries by edLMM

**Overview**. edLMM identified many significant genes associated with one of the seven diseases in the WTCCC dataset and two phenotypes in the NFBC dataset. By conducting literature search of these genes, we were able to annotate the functional relevance between significant genes and related diseases. For WTCCC data set, we chose Type 1 Diabetes (T1D) and Rheumatoid Arthritis (RA) for literature investigation as they reported the largest numbers of significant genes among the seven diseases. For each disease, we reported annotations of genes identified by edLMM but not EMMAX, two with the highest gene-disease association scores in DisGeNET and two novel genes not reported by DisGeNET(Pinero et al. 2015). (**T1D: Supplementary Tables S9; RA: Supplementary Tables S11**). For NFBC dataset, *NLRC5* from HDL is the only gene not reported by DisGeNET, and the other variants identified by edLMM but not EMMAX cannot be mapped to any genes using cS2G(Gazal et al. 2022). Therefore, we mainly focus on searching literature supports for *NLRC5* (**HDL: Supplementary Tables S15; LDL: Supplementary Tables S16**). The genes for which we have conducted literature search are listed in the **Supplementary Table S17** below:

**Supplementary Table S17**: Description of the genes for literature search in the WTCCC and NFBC datasets.

| Gene Name | Disease | p-value by EMMAX | p-value by edLMM | cS2G-UKBB-score | DisGeNET score |
|---|---|---|---|---|---|
| *PTPN22* | T1D | Not significant | 1.2000E-07 | 0.922 | 0.9 |
| *HLA-DRB1* | T1D | Not significant | 7.5700E-09 | 1 | 0.5 |
| *PANK3* | T1D | Not significant | 1.0300E-07 | 1 | Not reported |
| *BCL2L11* | T1D | Not significant | 3.5400E-08 | 1 | Not reported |
| *HLA-DQA2* | RA | Not significant | 3.2700E-12 | 1 | 0.4 |
| *PRRT1* | RA | Not significant | 5.1100E-08 | 1 | 0.1 |
| *SMOC2* | RA | Not significant | 3.4500E-08 | 1 | Not reported |
| *SCIMP* | RA | Not significant | 9.6600E-08 | 1 | Not reported |
| *NLRC5* | HDL | 6.5566E-08 | 1.4063E-08 | 0.655 | Not reported |

#### Type 1 Diabetes (T1D)

[*PTPN22*]: *PTPN22* encodes lymphoid-specific tyrosine phosphatase (LYP), an inhibitor of T cell activation. Many studies have revealed that *PTPN22* correlates strongly with the incidence of Type 1 Diabetes (T1D) and other autoimmune disease including Rheumatoid Arthritis (RA)(Bottini et al. 2006; Sarmiento et al. 2015). Particularly, *PTPN22* C1858T polymorphism has been confirmed associated with T1D in populations of Greek children and adolescents(Giza et al. 2013), and Caucasian descent from north central Florida(Zheng and She 2005) because of the increased frequency of 1858T allele in patients than in controls. Additionally, the risk contributed by this 1858T allele increased in patients with a family history of other autoimmune diseases, further supporting a general role for this variant on autoimmunity(Chelala et al. 2007).

Moreover, the *PTPN22* gene mutations played an important role in both Type 1 Diabetes (T1D) and Crohn's disease (CD) by modulating intracellular signaling(Sharp et al. 2015).

[*HLA-DRB1*]: *HLA-DRB1*, like another member (*HLA- DQB1*), has been found to hold many genetic determinants of type 1 diabetes(Erlich et al. 2008) and was regarded as one of the critical genotyping of markers for Type 1 diabetes(Nejentsev et al. 1999). According to studies, *HLA-DR* and *DQ* genes accounted for up to 50% of the familiar clustering of the type 1 diabetes(Rewers et al. 2003). Bedsides, this marker genes is not sensible to populations and has been reported to be associated with T1D in many populations across the world(Erlich et al. 2008; Mbanya et al. 2001; Sayad et al. 2013).

[*PANK3*]: *PANK3* belongs Pantothenate kinase (PANK) family, which catalyzes the rate-controlling step in coenzyme A (CoA) biosynthesis. *PANK3* Inhibitors have been widely used in the treatment of Type 2 diabetes(Leonardi et al. 2010). In additional, many studies have discovered the critical role played by PANK family genes in diabetes pathogenesis(Xiang et al. 2007; Zhang et al. 2007). However, the role played by PANK family genes in T1D still need to be explored future.

[*BCL2L11*]: *BCL2L11* was found to be one of the top five upregulated genes in T1D and its over expression might lead to the cell death of the islet β-cells in the pathogenesis of T1D(Fang et al. 2017). Moreover, recent studies suggest *BCL2L11* might function as a molecular biomarker for diabetic atherosclerotic plaque(Li et al. 2021) further suggesting the critical role played by *BCL2L11* in diabetes pathogenesis.

**Rheumatoid Arthritis (RA)**

[*HLA-DQA2*]: *HLA-DQA2*, together with other genes from HLA family (*HLA–DQB1*, *HLA–DRA*), contains many genetic determinants for RA, which have most reproducible associations across different ethnic groups(Negi et al. 2013). Many studies also reported that up-regulated *HLA-DQA2* is a common mechanism of RA(Pathi et al. 2021; Tan et al. 2017).

[*PRRT1*] In a prior extensive Mendelian randomization investigation that examined SNPs of multiple genes linked to IgG N-glycosylation, the only SNP identified as being related with RA was rs9296009. This SNP is present in the *PRRT1* locus, in chromosome 6 at positions 6p21.1–21.3 and in linkage disequilibrium with two other top RA SNPs, rs660895 and rs6910071(Wysocki et al. 2020). In another study, rs204999 (p-value 5.5x10-134), locating 6.2kb away from the 3′ end of *PRRT1*, is reported as one of the most significant SNPs associated with rheumatoid arthritis and juvenile idiopathic arthritis(Jia et al. 2020). Besides, *PRRT1* is also an eQTL gene that linked to top SNPs reported in autoimmune diseases(Liu et al. 2021). Although not much studies have explored the role played by *PRRT1* in Rheumatoid Arthritis, a lot of studies reported many differentially methylated regions in *PRRT* genes in many brain diseases. Moreover, multiple neurological illnesses in humans have been linked to mutations in *PRRT* family genes as well(Kim et al. 2022) (Ladd-Acosta et al. 2014).

[*SMOC2*] *SMOC2*, encoding a member of the secreted protein acidic, locates on chromosome number six in the 6q27 region. It is situated near the insulin-dependent diabetes mellitus 8 (*IDDM8*) locus, a genetic locus that is associated with autoimmune diseases such as rheumatoid arthritis and type I diabetes(Myerscough et al. 2000). Moreover, one recent research on the *SMOC2* gene showed that *SMOC2* (particularly, rs13208776 locating in *SMOC2*'s intron 4) is a significant candidate gene for susceptibility to generalized autoimmune vitiligo and maybe other autoimmune disorders(Birlea et al. 2010). Other studies on miRNAs demonstrated dysregulated miRNAs have an impact on several genes, including *ROR2, ABI3BP, SMOC2*, etc. in RA(Song et al. 2015).

[*SCIMP*] *SCIMP*, encoding an immune-restricted transmembrane adaptor protein, is a potential disease-related gene in human autoimmune disease(Lambert et al. 2013) (Dozmorov et al. 2014; Luo et al. 2017). Another recent work demonstrated *SCIMP* plays a crucial role in pro-inflammatory responses in macrophages because *SCIMP*-deficient mice display defects in pro-inflammatory cytokine production(Lucas et al. 2021).

**High-density lipoprotein (HDL)**

[*NLRC5*] *NLRC5*, also known as *NOD4* or *NOD27*, makes up a significant fraction and is involved in several immune responses. Recent studies have showed strong association between *NLRC5* and HDL(Hebbar et al. 2017; Hosseinzadeh et al. 2019). It should be highlighted that mounting evidence supports the involvement of *NLRC5* in innate immunity and inflammatory disorders. Furthermore, it has been demonstrated that *NLRC5* plays a crucial part in the regulation of a variety of signalling pathways. As a result of its extensive involvement in the emergence and progression of immunological disorders, *NLRC5* may be used as a possible therapeutic target for many chronical disease such as HDL(Wang et al. 2019). In addition, it has been reported that *NLRC5* deficiency leads to increased proliferation and migration of human aortic smooth muscle cells(Luan et al. 2019). Such smooth muscle cells from humans have been shown to express a variety of cholesterol uptake receptors, including the LDL receptor(Doran et al. 2008). With more smooth muscle cells available, LDL is likely to be transformed to HDL in large extent. Thus, deficiency of *NLRC5*, which cause proliferation of smooth muscle cells, might lead to increment of HDL density.
